# Sound comparison of seven TMS coils at matched stimulation strength

**DOI:** 10.1101/606418

**Authors:** Lari M. Koponen, Stefan M. Goetz, Debara L. Tucci, Angel V. Peterchev

**Author notes:** **Correspondence** Angel V. Peterchev, 40 Duke Medicine Circle, Box 3620 DUMC, Durham, NC 27710, USA.

## Abstract

**Background:** Accurate data on the sound emitted by transcranial magnetic stimulation (TMS) coils is lacking.

**Methods:** We recorded the sound waveforms of seven coils with high bandwidth. We estimated the neural stimulation strength by measuring the induced electric field and applying a strength–duration model to account for different waveforms.

**Results:** Across coils, at maximum stimulator output and 25 cm distance, the sound pressure level (SPL) was 98–125 dB(Z) per pulse and 75–97 dB(A) for a 15 Hz pulse train. At 5 cm distance, these values were estimated to increase to 112–139 dB(Z) and 89–111 dB(A), respectively.

**Conclusions:** The coils’ sound was below, but near, relevant exposure limits for operators and may exceed some limits for the subject. Exposure standards may inadequately capture some risks to hearing. For persons near operating TMS coils we recommend hearing protection, and we consider it essential for the TMS subject.

**Highlights:**

1. Coil click varies by over 20 dB(Z) between TMS coils at matched stimulation strength.
2. Close to TMS coil, sound pressure level may reach nearly 140 dB(Z).
3. For rTMS, the continuous sound level can exceed 110 dB(A).
4. Hearing protection is recommended during TMS, especially for the subject.

## Introduction

Transcranial magnetic stimulation (TMS) activates cortical neurons by electromagnetically inducing an electric field (E-field) pulse with a coil placed on the subject’s scalp. In addition to the desired E-field stimulation, the pulse causes mechanical vibrations in the coil, producing a brief but very loud sound impulse—often referred to as TMS coil click [1–2].

The sound pulse can potentially cause hearing loss. This risk can be mitigated with adequate hearing protection [3–5]. However, should the hearing protection fail, TMS can cause permanent hearing loss [6]. The sound also evokes unwanted neural activation that deteriorates the specificity of TMS and confounds functional brain imaging data [7], which can be only partially mitigated with noise masking [8].

Due to these side effects, the TMS safety consensus group suggested in 2009 that “the acoustic output of newly developed coils should be evaluated and hearing safety studies should be conducted as indicated by these measures” [9]. Since then, specific stimulator–coil combinations have been evaluated with hearing safety measurements [4–5], and a comparison between coils has been published [10]. To address methodological limitations of the latter, namely sound level measurements intended only for continuous (i.e., not impulsive) sounds based on IEC 61672 [2], and to expand the range of compared devices, we conducted measurements of the coil click in a total of seven coils for three stimulators and normalized the stimulation strength across the various configurations. Moreover, we interpret the results for single pulses as well as for repetitive TMS (rTMS) in the context of various standards for sound exposure limits.

## Material and methods

### TMS devices

We tested three stimulators with a total of seven different figure-of-eight coils. The model information of the tested stimulators and coils is provided in Table 1. The “stimulator output” refers to the stimulator capacitor voltage, expressed as a percentage of its maximum (maximum stimulator output, MSO), which determines the TMS pulse amplitude.

**Table 1.** Tested TMS stimulators and coils from Magstim (UK), MagVenture (Denmark), and Neuronetics (USA).

| Stimulator | Coil |
| --- | --- |
| Magstim Rapid2<br>(P/N 3013-US) | 70mm Double Coil (P/N 9925-00)* |
|  | AirFilm Coil (P/N 3910-00) |
| MagVenture MagPro X100 incl. MagOption<br>(P/N 9016E0731)** | C-B60 (P/N 9016E0482) |
|  | Cool-B65 (P/N 9016E0491) |
|  | MC-B70 (P/N 9016E0564) |
|  | MRi-B91 (P/N 9016E0661) |
| Neuronetics NeuroStar TMS Therapy System<br>(ver. 1.0, P/N 81-00315-000) | NeuroStar (ver. 1.0, P/N 81-00900-000) |

\* Connected to stimulator with adapter (Magstim, P/N 3110-00).
\*\* “Standard biphasic” pulse mode was used.

### Sound recording

A microphone was pointed at the center of the head-side of each coil at a distance of 25 cm (see Supplementary Fig. S1B). We sampled 10 pulses per stimulator output at 10% to 100% of MSO in 10%-MSO steps. The pulses at each stimulator output were separated by an interstimulus interval of 2–3 s. To isolate the coil click from sound originating from the stimulator, the coils were placed inside a soundproof chamber (Model 402.a; Industrial Acoustic Company, USA). To suppress echoes in the recordings, the relevant chamber interior walls were covered with 50 mm thick open-cell foam pads. To convert the digital signal into pascals, the system was calibrated with a 1 kHz, 1 Pa reference sound pressure source (Extech Instruments 407722, Extech, USA). To allow accurate measurement, we used an omnidirectional pressure microphone (Earthworks M50, Earthworks Audio, USA), which has a flat frequency response (–1.1 to +0.2 dB referenced to our 1 kHz calibration tone) from 3 Hz to 50 kHz and is free of distortion for sound pressure level (SPL) up to 140 dB(Z). The microphone was connected to a high-input-level-capacity preamplifier (RNP8380, FMR Audio, USA) and an audio interface with a sample rate of 192 kHz (Behringer U-Phoria UMC404HD, Behringer, Germany).

### Sound processing

In processing the raw audio, the first pulse at each stimulator output was excluded as an outlier, since it deviated from the subsequent pulses for all devices especially at low stimulator outputs. The signal was low-pass filtered at 60 kHz to remove high-frequency readout noise, and high-pass filtered at 60 Hz to remove low-frequency sounds which mostly originated from the air conditioning. The cut-off frequency for the latter filter was selected post-hoc, as none of the coils produced sound that deviated from the background noise at any frequency below 100 Hz. The high- and low-pass filters were fourth-order Butterworth type applied in both forward and reverse time, resulting in flat frequency response from 80 Hz to 50 kHz (within 1 dB). For these data, we computed the peak SPL [11] in a 0.2 s window around the pulse, the duration of the sound impulse defined as the period for which the sound pressure was within 20 dB of its peak value, and 1/3-octave spectra with the `poctave` function (Matlab R2018a, Mathworks, USA).

In addition, to estimate the continuous sound level due to rTMS, we synthesized 10 seconds of rTMS sound as a superposition of individual pulses repeated at 15 Hz. This frequency was chosen as the highest average frequency that is FDA-cleared for clinical use with the figure-of-eight type coils that we characterized, since the 15 Hz rTMS trains are equally loud to theta burst stimulation comprising bursts of three pulses at 50 Hz, repeated at 5 Hz [12].

### Estimation of stimulation strength

To compare the sound between different coils, which provided different stimulation waveforms and MSO, we normalized their stimulation strengths as follows. First, we measured the induced electric field (E-field) corresponding to a depth of 15 mm under the center of the coil in an 85 mm sphere approximating the head [13] using a 70-millimeter-high triangle-loop probe [14–15]. The E-field was sampled at 100 MHz with an oscilloscope (Tektronix MDO3054, Tektronix, USA). To obtain the effective stimulation strength, the E-field waveform was digitally low-pass filtered with a time constant of 200 μs, corresponding to the approximate strength–duration time constant in primary motor cortex [16–18]. The stimulation strength, defined as the absolute peak of the filtered waveform, was then scaled relative to the average resting motor threshold (RMT) by denoting 100% RMT to correspond to 50.3% MSO for the Magstim 70mm Double Coil driven by a Magstim Rapid biphasic stimulator [19]. Based on the variance of RMT across individuals, for about 95% of subjects, the individual RMT should lie between 75–125% of this value [19].

### Sound spectrum weighting and exposure limits

Choosing an appropriate exposure limit for TMS sound is not straightforward. In addition to the common limit of 115 dB(A) time-averaged steady-state sound level for short daily exposure [20–23], we must also consider the impulsive nature of the sound, which is similar to a small explosion, e.g., shooting a firearm, in terms of duration and peak sound pressure [24]. Consequently, measurement protocols for such sounds should provide good reference for the measurement protocol of TMS sound. Several safety standards give an instantaneous SPL limit of 140 dB, but with either A-weighting [22], C-weighting [23], or Z-weighting [20–21,25] (for frequency-weighting specifications, see Supplementary Fig. S2). A- and C-weighing only limit the sound in the hearing range (as they are defined only up to 20 kHz). With these, the limits for potential near-ultrasound content has to be considered separately. However, the relevant standards do not contain such separate impulsive limits, but only limits for continuous sounds of various durations, varying between 75–110 dB at 20 kHz and 110–115 dB at 25 kHz or above [26]. To avoid the need for two separate metrics, we use the Z-weighted 140 dB peak SPL from MIL STD 1474E [25], which also imposes a constraint on the impulsive near-ultrasound content. We also considered the sound level of rTMS pulse trains using A-weighting (slow time weighting) per IEC 61672 [11] computed with `splMeter` from the Audio Toolbox (Matlab R2018a, Mathworks, USA).

## Results

### Coil electric field and stimulation strength

The peak E-field at 100% MSO varied between 140–260 V/m across the tested coils, and it had nearly linear relationship to the stimulator output setting for all coils (Fig. 1A). The pulse durations varied between 176–353 μs and were largely stable across the output range of the devices except for the NeuroStar iron-core coil, which had about 19% longer pulse at 10% MSO than at 100% MSO (Fig. 1B). Consequently, for all but the iron-core coil, the devices showed an approximately linear relationship between the simulator capacitor voltage and the estimated stimulation strength (Fig. 1C) (NeuroStar devices, however, have an output calibration feature that can compensate for this nonlinearity).

**Figure 1.**
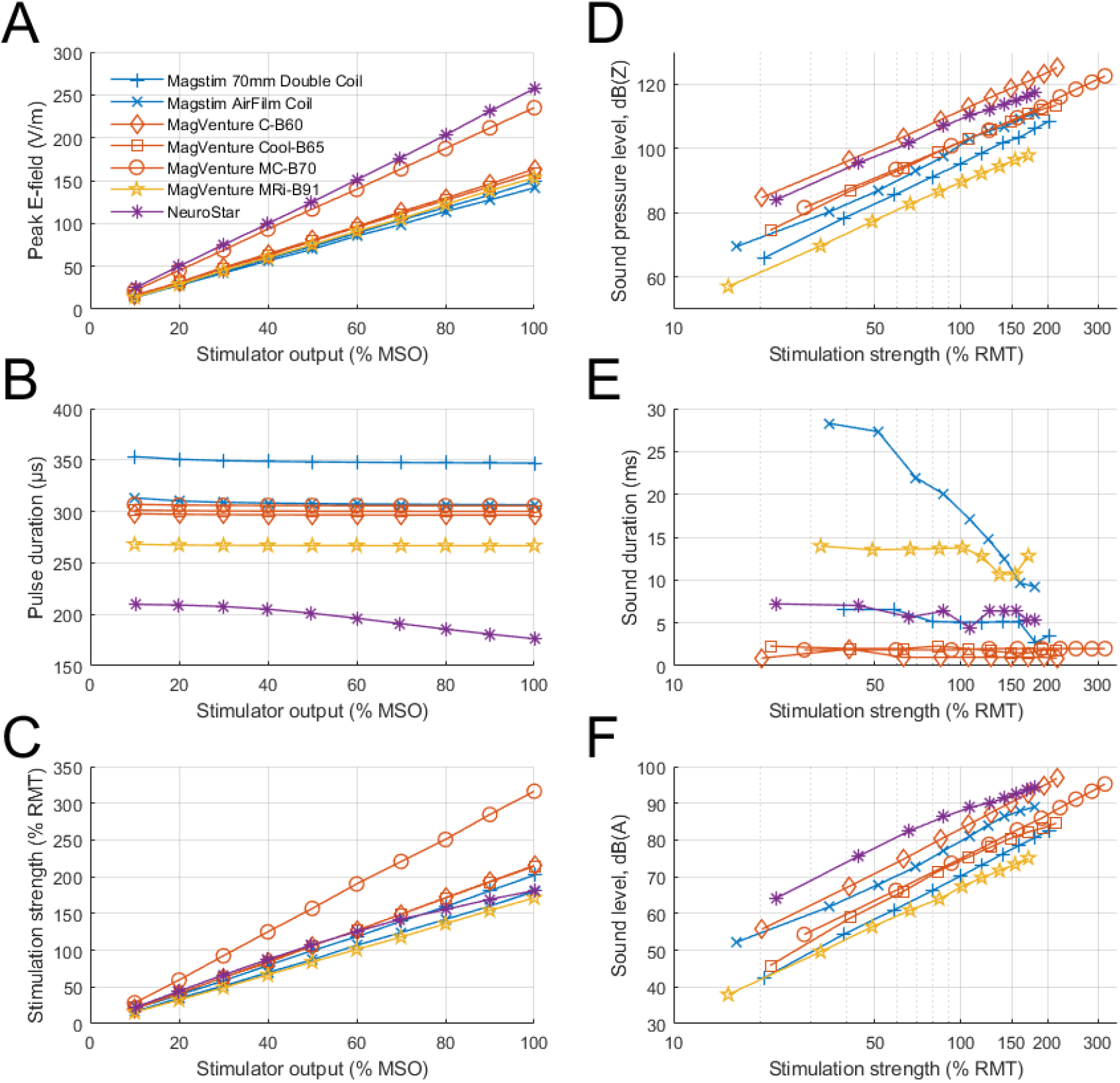
E-field and sound characteristics of various measured TMS coils. (A) Peak E-field value has approximately linear relationship with stimulator output. (B) Pulse durations of air-core coils are approximately constant, whereas pulse duration of NeuroStar iron-core coil starts to drop after about 30% MSO. (C) Stimulation strength estimated with neural model, where 50.3% MSO of Magstim 70mm Double Coil is set to correspond to 100% RMT. (D) Peak sound pressure level (SPL) with Z-weighting, i.e., flat response curve. (E) Duration for which sound is within 20 dB of peak value. (F) Simulated sound level of a 15 Hz rTMS pulse train with A-weighting (slow time weighting).

### Coil sound characteristics

At a distance of 25 cm, the seven coils had a wide range of peak SPL, 98–125 dB(Z) at 100% MSO and 92–114 dB(Z) at the estimated 120% RMT, with the MagVenture MRi-B91 being the quietest and the MagVenture C-B60 being the loudest (Fig. 1D). There were three distinct click types: very brief and sharp (1–2 ms) from conventional MagVenture coils; medium length (6–7 ms) from NeuroStar coil and Magstim 70mm Double Coil, and long (> 10 ms) from the MagVenture MRi-B91 coil and Magstim AirFilm Coil which have double casing (Fig. 1E). For most coils, the sound duration was comparable across stimulation strengths. For Magstim AirFilm Coil, the click duration decreased from 35 ms to 9 ms when the stimulator output was increased from 20% MSO to 100% MSO. All coils produced broad sound spectra with peak varying between 1–8 kHz and spectrum extending up to 30 kHz (Fig. 2). At 100% MSO, the 1/3-octave band had sound level between 40–69 dB at 20 kHz and 41–64 dB at 25 kHz. Peak SPL varied approximately proportionally to the stimulation strength squared (Fig. 1D). For example, increasing the stimulation strength by 20% increases the sound level by approximately 3 dB. This is expected from first principles, since the mechanical force causing the sound is a product of the magnetic field and coil current, both of which are proportional to the stimulator capacitor voltage.

**Figure 2.**
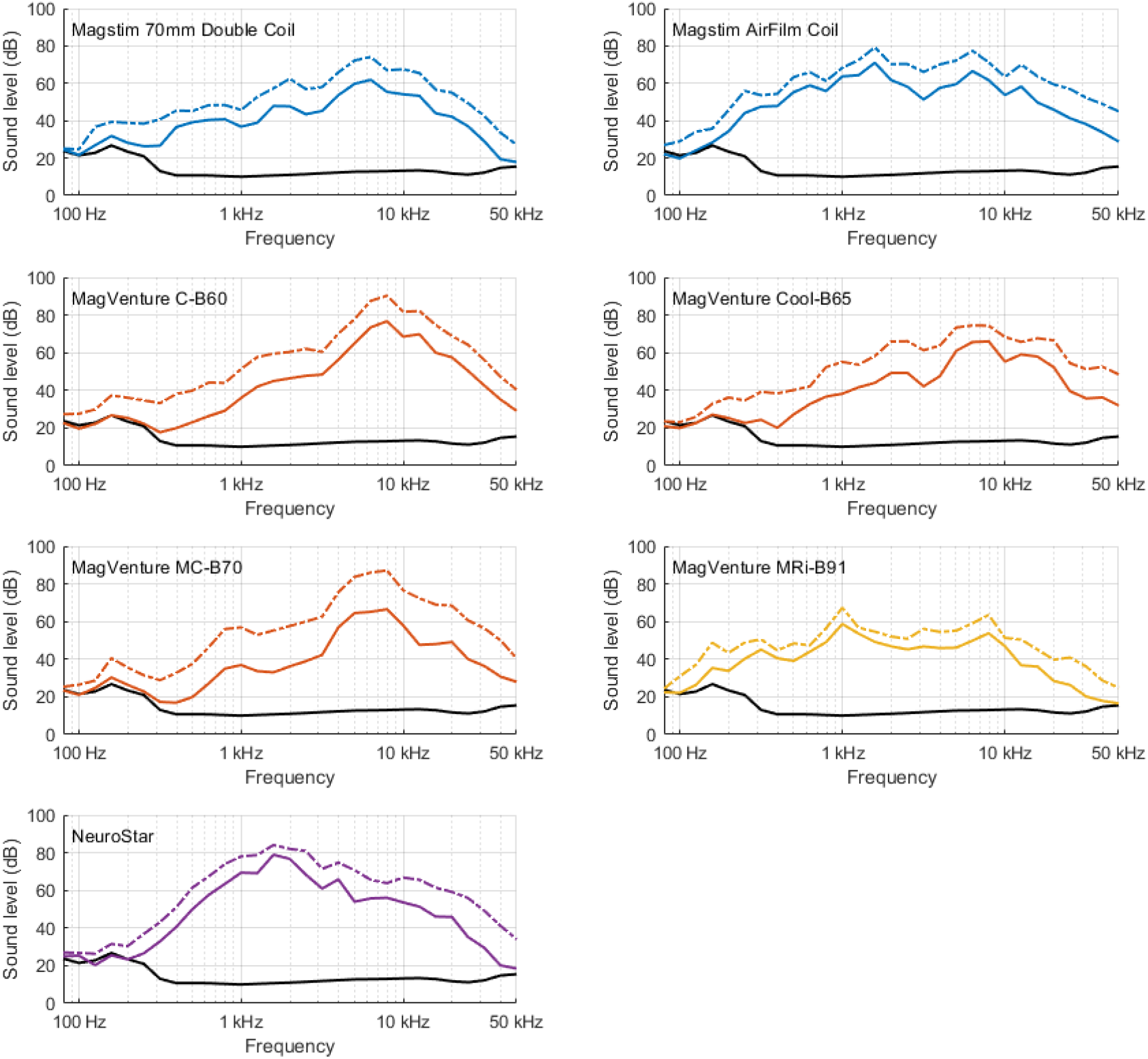
1/3-octave spectra of tested coils at estimated 100% RMT (solid line) and at 100% MSO (dash-dotted line). Solid black line shows ambient sound spectrum. For all coils, spectra power peak is between 1,000 and 10,000 Hz, i.e., within hearing range (20–20,000 Hz). Spectra at 100% RMT were produced from closest measured stimulator output by assuming that sound waveform is proportional to pulse energy. Due to the 0.2 s time window for each sample, the sound level corresponds to that of a 5 Hz rTMS pulse train.

The continuous sound for simulated 15 Hz rTMS at 120% RMT ranged 70–90 dB(A), with NeuroStar being the loudest and MagVenture MRi-B91 remaining the quietest but with a much smaller margin than for peak SPL (Fig. 1F). Doubling the repetition rate increases the sound level by approximately 3 dB; this scaling is valid for frequencies up to about 100 Hz, where the sound levels are 8 dB higher than the values shown in Fig. 1F. Table 2 summarizes how the data presented in Fig. 1 can be scaled approximately to various stimulation strengths, rTMS train frequencies, and coil distances.

**Table 2.** Scaling laws for reported peak SPL and rTMS sound level.

| Effect | Change in peak SPL | Change in rTMS sound level |
| --- | --- | --- |
| Stimulation strength | $12 \text{ dB} \cdot \log_2 \frac{\text{Stim. strength [\% RMT]}}{100\%}$ | $12 \text{ dB} \cdot \log_2 \frac{\text{Stim. strength [\% RMT]}}{100\%}$ |
| Repetition rate | 0 | $3 \text{ dB} \cdot \log_2 \frac{\text{Rate [Hz]}}{15 \text{ Hz}}$ |
| Distance from coil* | $-6 \text{ dB} \cdot \log_2 \frac{\text{Distance [cm]}}{25 \text{ cm}}$ | $-6 \text{ dB} \cdot \log_2 \frac{\text{Distance [cm]}}{25 \text{ cm}}$ |

\* Valid down to distance of 5 cm [27].

## Discussion

Our measurement setup mostly followed the suggested measurement procedures in MIL STD 1474E [25], which considers impulse sounds, such as from explosions, which are similar in their characteristics to the coil click. We measured the far-field sound, at a distance larger than the characteristic dimensions of the TMS coil. Our microphone was placed 25 cm away from the center of the coil, compared to coil–ear distances as small as 5 cm for the TMS subject. We chose the 25 cm distance to avoid either (1) having to sample the spatial distribution of the sound or (2) underestimating the SPL due to measuring near a nodal point of a standing wave pattern in the near field [2,18]. The sound pressure from TMS coils decays approximately inversely to the distance down to 5 cm [27] (see Table 2); therefore, the sound pressure at 5 cm from a coil is about 14 dB higher than the sound pressure measured at 25 cm. Thus, the peak SPL of the seven coils can be estimated to range from 106 to 129 dB(Z) at 120% RMT and 5 cm from the coils. The 22 dB difference between the coils is preserved with this scaling; this is comparable to the typical attenuation of air-conducted sound obtained with simple hearing protection [22]. Considering the extreme cases we can estimate, single pulses at 100% MSO and 5 cm from the coils, which for some TMS targets is comparable with the distance to the subject’s ear, produce SPL of 112–139 dB(Z).

As with the sound of single TMS pulses, there were large differences in the time-averaged steady-state sound levels of pulse trains for rTMS. Extrapolating to a 5 cm distance, a 15 Hz rTMS train has a sound level of 84–104 dB(A) at 120% RMT. At 100% MSO, the sound level would reach 89–111 dB(A). These values would be approximately 2 dB lower for 10 Hz rTMS trains. The OSHA limit for continuous sound level is 115 dB(A) for short durations (< 15 min) and degrades by 5 dB per doubling of exposure duration to 90 dB(A) for exposure of 8 hours [20–21]. Even in case of a TMS operator administering multiple rTMS sessions per day and remaining in close proximity of the coil at 25 cm, our estimated sound levels are below the OSHA limit for occupational exposure (with the possible exception of very high repetition rates, e.g. 100 Hz used in magnetic seizure therapy). However, these exposure limits are approached for TMS subjects. Moreover, for most coils the sound level extrapolated to a distance of 5 cm can exceed the lowest near-ultrasound limit of 75 dB at 20 kHz [26]. Therefore, the use of hearing protection for the TMS subject is advisable.

In this dataset of commercial stimulators and coils, all coils had most of their sound in the hearing range. All coils, however, also had considerable sound level in the 20–30 kHz range, with 15 Hz rTMS at 100% MSO exceeding the lowest near-ultrasound limit of 75 dB in the 1/3-octave band at 20 kHz [26] at 5 cm from the coil for all but Magstim 70mm Double Coil and MagVenture MRi-B91 (Fig. 2). Thus, to fully capture the sound, we recommend measuring the sound to at least 30 kHz, which is much higher than the 8 kHz or 16 kHz bandwidth available in common sound level meters compliant with Class II or Class I of IEC 61672 (even if the meter supports measurement of peak SPL in addition to the time-averaged steady-state sound level of this standard).

The previous comparison between TMS devices used sound level meters with a slow time weighting and a measurement bandwidth of only 8 kHz [10], which is insufficient for impulsive sounds [2]. Indeed, we obtained significantly higher sound pressure levels, and, for the three coils common in these two studies, a different order. At 100% MSO and 25 cm distance in our study, the Magstim 70mm Double Coil, MagVenture Cool-B65, and NeuroStar coils registered at 108, 113, and 117 dB(Z), respectively, whereas these coils produced only 90.2, 78.3, and 82.7 dB(A), respectively, in the prior study despite the smaller measurement distance of 10 cm [10]. Specifically, the Magstim and MagVenture coils were respectively 18.3 dB and 35.1 dB louder with the wide-bandwidth impulse sound measurement at 25 cm, and these numbers are estimated to rise to 26.2 dB and 43.1 dB extrapolating the loudness at a matched distance of 10 cm. It should be noted, though, that the version of a particular coil that we tested may have differed from those in other studies; for example, the NeuroStar coil we tested is an early model (version 1.0).

On the other hand, compared to the previous peak SPL measurement using a head phantom [5], we obtained lower SPLs for the same coil (Magstim 70mm Double Coil, 108 dB(Z) in free field versus 127.6 dB(C) in the ear canal). This difference is only partially explained by the smaller distance between the ear and the coil. The main factor is likely the ear canal model itself, which mimics the fact that the pressure levels in the ear are higher than the free field pressure levels [29]. The GRAS RA0045 microphone used in those measurements attempts to mimic human ear-canal response curve between 0 and 10,000 Hz (and has a narrow +32 dB resonant peak outside that measurement range at 13.5 kHz, likely amplifying the high-frequency content of the coil click) [28]. All mentioned 140 dB limits, however, are defined for free-field SPL [20–23,25]. Importantly, head phantoms established in acoustics do not model body sound through the skull, which depends on the stimulation location and can increase the relevant sound pressure of TMS in the ear [30].

Our measurements indicate that for an operator holding a TMS coil, the sound from single-pulse TMS and rTMS at common frequencies is below relevant exposure limits. However, the sound level can come close to the exposure limits, or even exceed them at very high pulse repetition rates. Moreover, there is emerging evidence that hearing loss is more complex than assumed in the current exposure limits [31]. So called “hidden” hearing loss, i.e., a reduction in the ability to distinguish sounds in noisy environment, has been observed already with loud noise exposure below levels causing permanent audiometric hearing loss [31]. Further, the standard limits [20–21,23,26] are only a guide to maximum allowable environmental exposure, and do not guarantee safety [21,32] (i.e., one should reduce exposure well below the specified limits, if possible). Finally, we characterized only a sample of figure-of-eight TMS coils; it is possible that there are or will be louder ones. Indeed, rTMS with a different kind of coil has caused permanent hearing loss [6]. Based on these considerations, we recommended that persons close to an operating TMS coil wear hearing protection.

For the TMS subject, the sound levels are even higher. Further, conventional acoustic measurements of TMS coils do not address the sound conducted through the skull, neither do they account for the possibility of significant near-field peaks of acoustic energy. Therefore, we consider hearing protection essential for TMS subjects.

## Conclusion

Commonly used TMS devices have large differences in their sound levels at matched stimulation strength. The sound of the tested coils was below, but near, relevant exposure limits at distances relevant for the TMS operator. For the TMS subject, some near-ultrasound limits may be exceeded and there may be other sources of risk such as skull conduction and near-field sound pressure peaks. Generally, sound exposure standards may inadequately capture some risks to hearing. Therefore, we recommend hearing protection for persons near operating TMS coils, and we consider hearing protection essential for the TMS subject.

## Supporting information

Sound recordings

Electric field recordings

Probe drawings

## Acknowledgements

Research reported in this publication was supported by the National Institute of Mental Health of the National Institutes of Health under Award Number R01MH111865. The content is solely the responsibility of the authors and does not necessarily represent the official views of the National Institutes of Health.

## Supplementary methods

### Sound recordings

Based on preliminary measurements, the impulsive sound from TMS pulse generators can be relatively loud. In preliminary measurements with the Tonica MagPro MST stimulator (MagVenture, Denmark), using the same 25 and 100 cm distances from the coil to the microphones as in the final measurement dataset, an impulsive sound source with a peak SPL of 104 dB(Z) at 1.7 meters away from the farther microphone was identified. This distance corresponded with the distance to the stimulator. This sound was just 9 dB less than the sound from the tested coil at 1.0 m, i.e., only about 6 dB less if both sound sources would be at equal distances.

To measure accurately the waveform of the coil sound, we had to isolate it from the impulsive sound of the stimulator. We achieved this isolation by measuring the coils inside a soundproof chamber with the TMS pulse generator outside. If one is interested only in the peak SPL and measures from a reasonably close distance to the coil, a soundproof chamber will not be needed. However, in such case, one might want to use at least two microphones (and beamforming) to identify and, if needed, to reject the sound not originating from the direction of the coil. The supplementary sound recording dataset contains a second microphone at 100 cm from the coil, but due to the soundproof chamber the data from this microphone was not needed in the analysis. For the placement of the microphones and the acoustic foam, see Fig. S1B.

The peak SPL was defined as

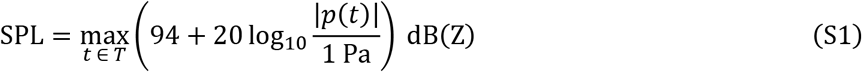

where *T* is the 0.2 s time window around a trigger event (from –50 ms to 150 ms) and *p*(*t*) is the filtered instantaneous sound pressure [S1]. The ambient noise measured in the soundproof chamber had SPL of 45.0 dB(Z) (95% confidence interval of 43.3–46.6 dB(Z)) with Z-weighting and sound level of 21.2 dB(A) with A-weighting. Consequently, we could measure the pulse durations for pulses down to about 70 dB(Z), i.e., down to 20% MSO for the three quietest coils and down to 10% MSO for the other coils.

As a supplementary material we provide the full set of sound recordings, band-pass filtered from 60 to 60,000 Hz, as described in the methods, and compressed to contain data from –50 ms to 150 ms relative to each pulse, in Matlab file “sound.mat”. In this file, the originally 24-bit sound recording data were converted to Pascals and stored as 32-bit floating point numbers. In addition to the coil sounds, to assess the background noise level, there are also recordings of 100 events in which no pulse was delivered.

### Electric field measurements

The coil E-field was measured with a 70-millimeter-high printed-circuit-board triangular pickup loop with a 50-ohm series resistor, a 3d-printed holder, and a laser-cut alignment tool for mounting the holder on the coils (Fig. S1A). The E-field measurements were estimated to have < 2 mm positional (< 0.05 mm for distance from the bottom of the coil) and < 2° angular uncertainty. With these uncertainties, the field values for each coil have < 2% gain uncertainty. The technical drawings of the measurement probe are provided in file “probe.zip” containing the Gerber files for the two-layer printed-circuit-board probe, an STL file for the 3d-printed probe holder, and a SVG drawing of the laser-cut alignment plate.

The full E-field measurement dataset, including the pulse waveforms, with one pulse per coil per stimulator output, sampled at 100 MHz, is provided in Matlab file “stimulation.mat”.

The stimulation strength was computed from the E-field recordings as

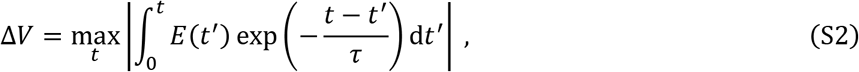

where Δ*V* is the stimulation strength, *E*(*t*) is the measured E-field waveform, and *τ* is the strength–duration time constant, here 200 μs.

---

**Figure S1.**
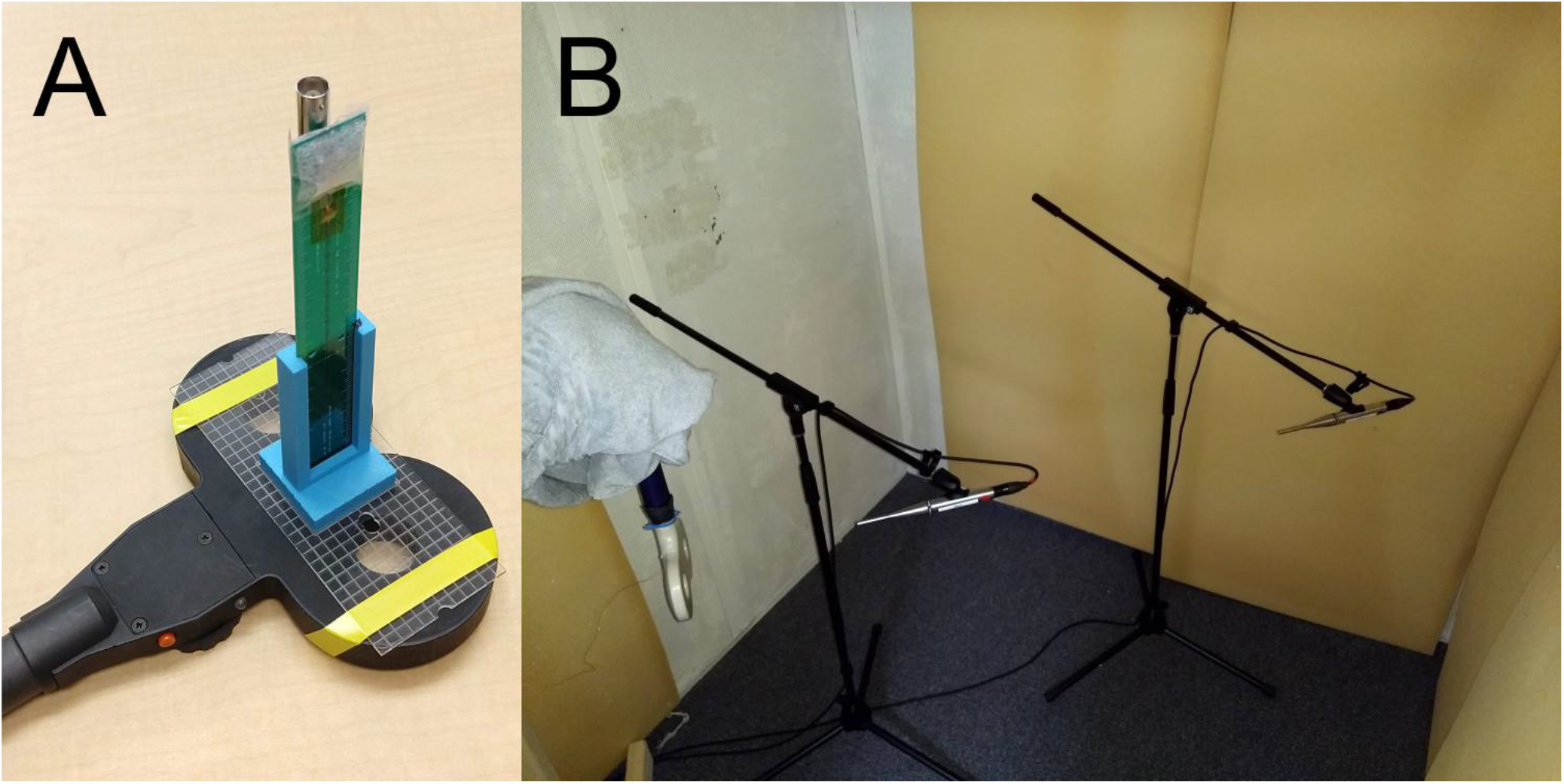
(A) Triangular E-field probe in 3d-printed holder on laser-cut alignment plate positioned on example TMS coil. (B) Sound-recording setup: two microphones are directed at head-side of coil at respective distance of 25 cm and 100 cm measured from microphone capsule. Soundproof chamber provided adequate attenuation of external sound sources including TMS pulse generator; therefore, only data from closer microphone was ultimately needed for analysis. Corners behind coil and microphones were covered with 50 mm thick open-cell foam to suppress echoes. Coil cable is wrapped in towels to suppress its sound.

**Figure S2.**
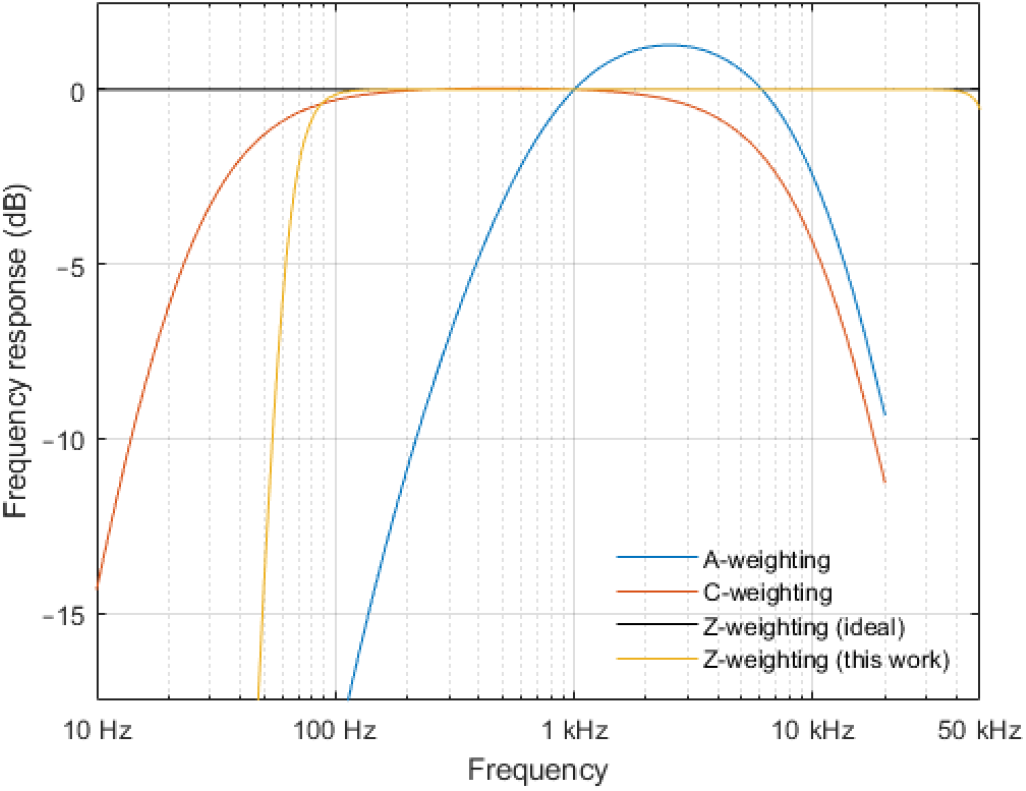
Frequency weightings used in hearing safety standards. A-weighting roughly mimics perceived loudness curve for barely audible sounds, whereas C-weighting mimics that of loud sounds. Ideal Z-weighting corresponds to no weighting at all. In this work, Z-weighting filter had passband from 80 to 50,000 Hz, with sharp low-frequency cut-off selected to reduce noise from non-TMS-coil sources. In practice, a Class II sound-level meter may omit anything above 8 kHz, and a Class I sound-level meter anything above 16 kHz.

